# Dissociating Goal from Outcome During Action Observation

**DOI:** 10.1101/2023.10.31.564940

**Authors:** Shuchen Liu, Moritz F. Wurm, Alfonso Caramazza

## Abstract

Understanding the goal of an observed action requires computing representations that are invariant to specific instantiations of the action. For example, we can accurately infer the goal of an action even when the agent’s desired outcome is not achieved. Observing actions consistently recruits a set of frontoparietal and posterior temporal regions, often labeled the "action observation network" (AON). While progress has been made in charting which regions of the AON, are involved in understanding goals of observed actions, it is not clear where goals are represented independently of outcomes. We used fMRI-based multivariate pattern analysis to identify such regions. Participants watched videos of successful and failed attempts of actions with different goals involving 2 different object types. We found that bilateral anterior inferior parietal lobe and right ventral premotor cortex distinguished between object-specific action goals regardless of outcomes. Left anterior inferior parietal lobe encodes action goals regardless of both outcomes and object types. Our results provide insights into the neural basis of representing action goals and the different roles of frontoparietal and posterior temporal regions in action understanding.

## Introduction

When observing other people’s behaviors, we can accurately interpret what they intend to do and evaluate whether they have achieved their desired result through their actions. Imagine a person reaching for a glass of water and knocks it off the table. We infer that they are likely trying to take and drink from the glass instead of breaking it. Even though many possible desires of the agent can lead to the same consequence, we can effortlessly understand what a person wants to do. This ability to accurately infer the agent’s goal allows us to communicate and interact with others effectively. What is the neural substrate that supports such ability? Where is action goal represented in the brain?

The goal of an action can be described at many levels (Hamilton & Grafton, 2006; Keele et al., 1990). For example, when someone is opening a bottle of water, their goal can be thought as “to open the bottle”, “to drink”, or “to quench thirst”. In this study, we restrict our search to the level that describes the immediate physical consequence on a target object, which an agent wishes to achieve through their motor action (i.e., “to open the bottle”). To identify regions that represent such action goals, the neural representation of a goal should be invariant to the specific motor implementations of the action and, crucially in the present context, the outcome of the action.

Neuroimaging studies have found that observing actions consistently recruits a set of frontoparietal and posterior temporal regions, including ventral premotor cortex (PMv), inferior parietal lobe (IPL), and lateral occipitotemporal cortex (LOTC) (e.g., Caspers et al., 2010; Kilner, 2011; Van Overwalle & Baetens, 2009; Urgesi et al., 2014; Wurm & Lingnau, 2015). These regions together are often labeled the “action observation network” and are thought to play important roles in understanding other people’s actions. Interpreting the goal of an action is a crucial part of action understanding, and it is natural to assume that action goals are represented somewhere in this network.

Several regions in the action observation network have been implicated to contribute to goal understanding, including left anterior intraparietal sulcus, right inferior frontal gyrus, right inferior parietal lobe and lateral occipitotemporal cortex (Hamilton & Grafton, 2006; Hamilton & Grafton, 2008; Lingnau & Downing, 2015; Ramsey & Hamilton, 2010; Wurm et al., 2016). However, there has been no clear convergence on what precise roles each region plays in action recognition and how they relate to representing action goals. Critically, these studies all used stimuli where the actions were complete and well-executed. In other words, the desired outcomes were always achieved, so the goal of an action was always conflated with the outcome. Thus, it is not clear whether it is the goals or the outcomes that were represented in these regions.

In the current study, we investigated, using functional magnetic resonance imaging, where in the brain action goal is represented independently of outcome. We scanned participants while they viewed videos of a human agent performing an action on an object. Each action had one of the four possible goals: “to open a bottle”, “to open a bag”, “to close a bottle”, or “to close a bag”. To the end of disentangling goal and outcome, each action could either be successfully executed or failed to achieve the desired outcome (Fig. 1A). That is, an action with a specific goal could end in different outcomes. We used multivoxel pattern analysis (MVPA) to identify brain regions that potentially encode action goal independently of outcome. Specifically, we predict that if a region represents action goal as such, its neural patterns should be more similar when activated by actions that share the same goal even if they have different outcomes than by actions with different goals. Some of the regions which were considered to represent action goal may in fact represent outcome. In such regions, we do not expect to observe effects of encoding goals independently of outcome.

**Figure 1.**
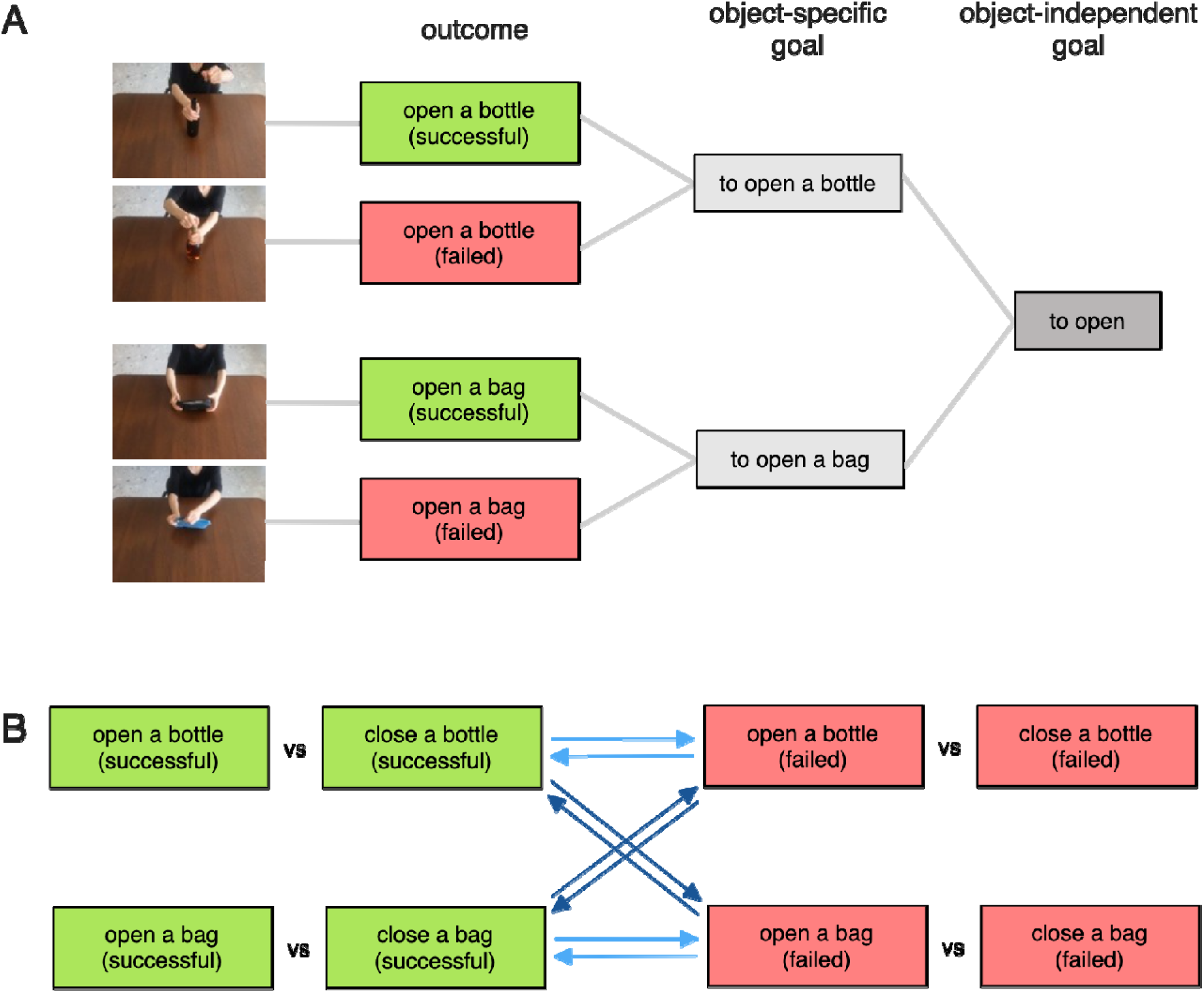
**(A)** Levels of goal representation. The left column shows specific instantiations of an action goal ending in different outcomes. The object-specific level (middle column, light gray) describes an action goal with a specific target object, generalized across different outcomes. The object-independent level (right columns, dark gray) describes an action goal that generalize across both outcomes and object types. **(B)** Schema of decoding actions across outcomes. Light-blue arrows: decoding across outcomes and within object types. Dark-blue arrows: decoding across outcomes and across object types

To control for the effect of perceptual information that are irrelevant to action goals, we included different perceptual variants for each unique goal, involving different moving directions, viewpoints, main hand, and object exemplars. In addition, by involving two different types of objects (i.e., bottles and zipper bags), we also investigated whether action goals are also represented independently of both outcome and the type of target object involved in the actions. That is, we explored which parts of the brain capture the similarity between the goal “to open a bottle” and the goal “to open a bag”.

We found that neural patterns in right anterior IPL and right PMv distinguished between object-specific action goals independently of outcomes. In left IPL, the neural patterns discriminate action goals independently of both outcomes and object types.

## Results

We scanned thirty-eight participants, thirty-two of whom were included in the analysis (see *Methods*). They watched the action videos and performed a catch trial detection task by pressing a button when the current action was neither opening nor closing. Participants performed well in the task, with a mean miss rate of 6.32% ± 6.66% (SD) and a mean false alarm rate of 1.84% ± 2.97% (SD).

### Decoding object-specific successful actions

We first aimed to identify regions that generally encode information about actions. To this end, we performed a within-object classification of successful actions in a whole-brain searchlight analysis (See *Methods* for details). We trained a classifier to discriminate the neural patterns activated by successful opening actions vs successfully closing actions involving the same object type. The results revealed a network of occipitotemporal and frontoparietal regions (Fig. 2A) that are well-aligned with the action observation network identified in previous studies (e.g., Caspers et al., 2010; Kilner, 2011; Van Overwalle & Baetens, 2009). We predicted that action goals are represented somewhere in this network, therefore we used the corrected map from this analysis as a mask to constrain the search space of following analyses.

**Figure 2.**
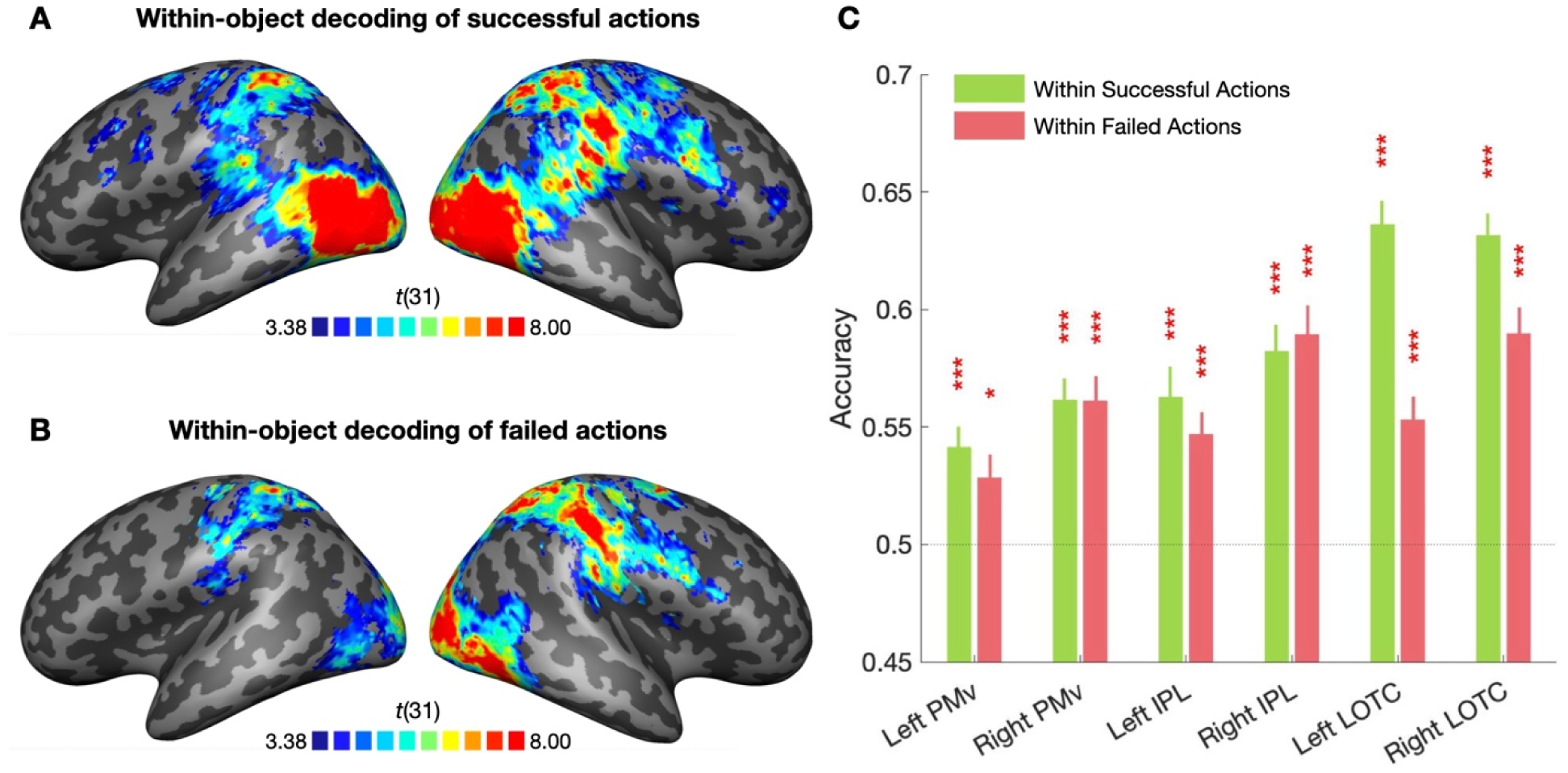
Within-object and within-outcome decoding of actions. Results of whole-brain searchlight for **(A)** within-object decoding of successful actions and **(B)** within-object decoding of failed actions (one-tailed t-test against chance 50%). Only clusters surviving Monte Carlo Cluster correction (p_initial_ = 0.001) are shown. The searchlight for (B) was conducted within the mask created from the map in (A); **(C)** ROI decoding accuracies of within-object decoding for successful actions and for failed actions, respectively. Error bar indicates SEM; asterisks indicate significance (FDR-corrected, * p < 0.05,** p < 0.01,*** p < 0.001)

In ROI analyses, we took a closer look at the core regions of the network, which were defined independently of the searchlight analyses. All ROIs, i.e., bilateral PMv, IPL and LOTC, showed significantly above-chance decoding accuracies (FDR-corrected, all ps < 0.0223, Fig 2C), confirming that these regions encode information about actions.

### Decoding object-specific failed actions

In effective action understanding, we should be able to infer the agent’s goal even if the action has failed and we do not observe the desired outcome. In previous studies that aimed to explore where goals are represented in the brain, participants were always observing actions that had succeeded. Thus, the effect could be driven by the outcomes instead of the goals. To disentangle these two aspects, we introduced failed action stimuli, where the agent did not achieve their goal and the object ended in the same state as it was before the action. We predict that if a region represents action goal per se, it should be differentially activated by actions with different goals even when the actions failed.

To test this, we performed a within-object classification of failed actions. We conducted a searchlight analysis within the network identified in the previous test. This analysis revealed postcentral gyrus, superior parietal lobule, occipital cortex in both hemispheres, and extended to ventral premotor cortex and posterior temporal lobe in the right hemisphere (Fig. 2B). All ROIs showed significantly above-chance accuracies, consistent with the searchlight results (FDR-corrected, all ps < 0.023, Fig. 2C). A three-way repeated-measures ANOVA with factors Decoding Tests, ROI, and Hemisphere showed a significant interaction of Decoding Tests x ROI (*F*(2, 62) = 14.96, *p* < 0.001) and an interaction of Decoding Tests x Hemisphere (*F*(1, 31) = 8.83, *p* = 0.006), as well as main effects for Decoding Tests (*F*(1, 31) = 8.67, *p* = 0.006), ROI (*F*(2, 62) = 39.74, *p* < 0.001), and Hemisphere (*F*(1, 31) = 31.72, *p* < 0.001). Post-hoc two-tailed paired t-tests revealed weaker decoding of failed actions comparing to decoding of successful actions in left LOTC (*t*(31) = 6.579, *p* < 0.001) and right LOTC (*t*(31) = 3.479, *p* = 0.0015), but not in other regions. We also observed a right lateralization effect: All three right ROIs showed stronger decoding of object-specific failed actions than their left-hemisphere counterparts (FDR-corrected, all ps < 0.001)

This result suggests that PMv, IPL and LOTC in both hemispheres could distinguish actions with different goals even when the desired outcomes of the actions were not observed. However, this does not mean that these regions are encoding action goals per se. In fact, the actions in the training data and the actions in the testing data shared the same outcomes. Therefore, it is not clear from just this result what information – action outcome or goal – is encoded in these regions.

### Decoding object-specific actions across outcomes

To identify regions that encode information about action goals independently of outcome at an object-specific level, we performed a within-object classification of action goals across outcome. That is, we trained the classifier on successful actions involving a specific object (e.g., successfully opening a bottle), and tested it on failed actions involving the same object (e.g., failed to open a bottle), and vice versa (Fig. 1B). These two actions involve the same goal of “to open the bottle”, but one ended in an opened bottle, the other ended in a closed bottle. Our reasoning is that if a region represents goals per se independently of outcome, their neural patterns should be similar when activated by actions with the same goals even if they have different outcomes. In other words, the representation of action goals in this region should generalize across different outcomes.

This searchlight analysis showed cross-outcome generalization of actions in left anterior IPL, right anterior IPL, and right PMv, corrected for multiple comparisons (Fig. 3A). We did not observe effects in LOTC in either hemisphere. A ROI analysis showed significantly above-chance accuracies in left IPL and right PMv (FDR corrected, ps < 0.005, Fig. 3C). However, we did not find above-chance decoding in our right IPL ROI. Notably, the right IPL ROI is more superior to the right IPL peak we identified in the searchlight analysis, which could explain the discrepancy between searchlight and ROI results.

**Figure 3.**
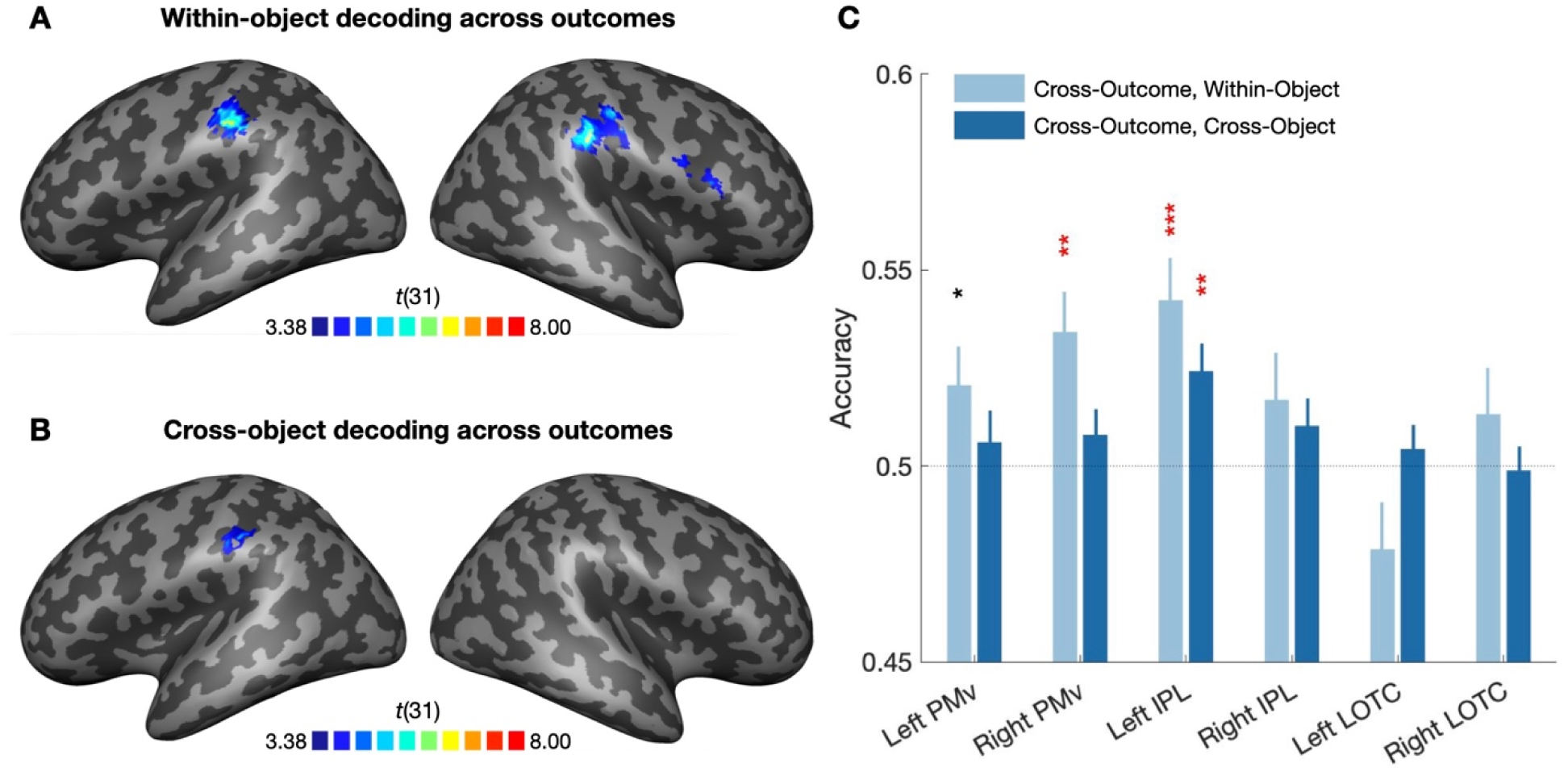
Decoding action goals across object types. **(A)** t map of within-object decoding of action goals across outcomes (one-tailed t-test against chance 50%). Only clusters surviving Monte Carlo Cluster correction (p_initial_ = 0.001) are shown; **(B)** t map of cross-object decoding of action goals across outcomes (one-tailed t-test against chance 50%), threshold at *p* < 0.001, uncorrected; **(C)** ROI decoding accuracies of decoding action goals across outcomes, for within-object and for cross-object, respectively; error bar indicates SEM; asterisks indicate significance (black: uncorrected, p <0.05; red: FDR-corrected, * p < 0.05,** p < 0.01,*** p < 0.001)

### Decoding actions across outcomes and across object types

In the preceding analysis, we identified regions that encode the goals independently of outcome in actions that involve the same object type. Do left aIPL, right aIPL, and right PMv encode the same information about the action goals? Previous studies have provided abundant evidence that different parts of AON may play different roles in action recognition (e.g., Wurm & Lingnau, 2015; Wurm et al., 2019b). For example, Wurm & Lingnau (2015) found that PMv discriminated different actions involving the same object, whereas in IPL and in LOTC, it was possible to decode actions across different object types (e.g., bottles and jars). However, it is not clear what information about these actions is encoded in these regions. Since two types of objects, bottles and zipper bags, were included in our stimuli, we conducted an exploratory analysis to examine whether action goal is represented independently of both outcome and of object type. This allowed us to explore whether the regions we found in the preceding analysis, namely, left aIPL, right aIPL, and right PMv, represent different information about action goals.

We performed a cross-outcome cross-object decoding (Fig. 1C). Specifically, we trained the classifier to discriminate between actions with a specific outcome and a specific object (e.g., successful bottle actions) and tested with actions with a different outcome and a different object (failed bag actions). After Monte Carlo Cluster correction, we did not find any significant cluster (p_initial_ = 0.001). However, we found one cluster in the left anterior IPL in the uncorrected map of one-tailed t-test of individual accuracies against chance (*p* < 0.001, Fig. 3B). This is consistent with the ROI results, which showed that the left IPL ROI had significantly above chance accuracy in cross-outcome cross-object decoding (FDR corrected, p < 0.001, Fig. 3C).

The other two regions revealed in the object-specific cross-outcome decoding, namely right IPL and right PMv, were absent here. To further compare the current results to the preceding results from within-object cross-outcome decoding, we performed a repeated-measures ANOVA with factors Decoding Test, ROI, and Hemisphere. Results showed significant interactions for Decoding Test x ROI x Hemisphere (*F*(2, 62) = 5.94, *p* = 0.004), Decoding Test x ROI (*F*(2, 62) = 4.05, *p* = 0.022), ROI x Hemisphere (*F*(2, 62) = 9.45, *p* = 0.003), as well as a main effect for ROI (*F*(2, 62) = 7.57, *p* = 0.011). Post-hoc two-tailed paired t-tests confirmed that right PMv showed significantly lower accuracy at decoding action goals across outcomes and objects than only across outcomes (*t*(31) = 2.212, p = 0.035). The right IPL ROI showed similar accuracies in these two decoding tests (*t*(31) = 0.487, p = 0.630). However, this was expected because the right IPL ROI was considerably superior to the peak found in the whole brain search. We also confirmed that the decoding accuracy was significantly higher in left IPL ROI than right IPL (*t*(31) = 2.192, p = 0.036) and right PMv (*t*(31) = 2.106, p = 0.043). This result suggests that left IPL, but not right IPL or right PMv, contains information about action goals both at an object-specific level and at an object-independent level.

## Discussion

We used MVPA to examine where action goals are represented independently of outcomes in the action observation network. The main findings were the following: (1) left anterior IPL, right anterior IPL and right ventral PM discriminated action goals independently of outcome at an object-specific level (i.e., involving the same object type); that is, these regions respond similarly to actions with the same goal and object type but different outcomes (e.g., “successfully opening a bottle” and “failed to open a bottle”) and respond differently to actions with different goals but the same outcome (e.g., “successfully opening a bottle” and “successfully to closing a bottle”); (2) neural patterns in left anterior IPL additionally discriminated actions with different goals independently of both outcome and object type; that is, this region responds similarly to the action “successfully opening a bottle” and “failed to open a bag”, even though they involve different object and end in different outcomes.

We found that action goals are represented independently of outcome in left aIPL, right aIPL and right PMv. Can these results merely reflect the effects of extraneous factors confounded with goals? One potential factor is kinematic feature similarity. For example, in both “successfully opened a bottle” and “failed to open a bottle”, the agent was trying to pull the cork out of a bottle. So, these two conditions were very similar in hand movements. We had anticipated this potential problem when creating the stimuli for the relevant comparisons and made the hand trajectories of opening and closing actions of the same object as similar as possible. Thus, it is not very likely that this generalization was purely driven by the nuanced differences between the simple kinematic features of opening and closing actions. Our finding suggests that these regions encode information about the agent’s desired physical consequence of object-directed actions, regardless of the actual physical consequence.

An important feature of our results is that right aIPL and right PMv encode information about action goals only across outcomes, but not across object types. This suggests that, for transitive object-directed actions, there is a level of representation such that the target object is a constitutive part of the goal. In other words, the representation of a goal is instantiated in a manner that includes the specification of a particular target object, such as, for example, “to open the bottle” is a goal, which is a different goal from “to open the bag”.

Interestingly, in comparison to right aIPL and right PMv, we were able to decode goals across both outcomes and object types in left aIPL. This dissociation between left aIPL and right aIPL/right PMv indicates that they may encode different information about action goals. We speculate that right aIPL and right PMv may represent information that are specific to the type of object involved, e.g., the motor components required to manipulate a particular object into the desired state. In contrast, left aIPL responds similarly to the action “successfully opening a bottle” and “failed to open a bag”. More intuitively, in left aIPL, the goal “to open a bottle” is considered more similar to “to open a bag” than to “to close a bottle”. We speculate that this region may represent information that are common across the goals to open different object types, for example, to “make the contents of a container accessible”.

Our findings are in line with other studies, which suggest a similar role of the inferior parietal cortex in representing aspects of action processing. Desmurget et al. (2009) found that stimulation in posterior parietal cortex led to patients having a desire to move, in the absence of actual movement responses. Patri et al. (2020) reported that tCBS disruption to a site in left anterior IPL did not affect participants’ performance in discriminating kinematic features of observed actions but impaired their ability to infer the goal of the action based on these kinematic features. These findings suggest that during action execution, IPL may serve as a transition hub for translating motor intents into actual motor movements, whereas in action observation, IPL may be involved in goal understanding by mapping kinematic features to the correct goal.

Our findings revealed an interesting dissociation between the frontoparietal regions and LOTC, further supporting the notion that different parts of the AON make different contributions to action recognition. In LOTC, the representations of actions could both discriminate successful actions and discriminate failed actions. However, they did not generalize across outcomes at neither the object-specific level nor the object-independent level. This dissociation suggests that the frontoparietal regions represent the inferred or “invisible” goal, whereas LOTC seems to represent what actually happened in the observed action. Importantly, the effect of decoding actions with different goals in LOTC was stronger for successful actions than failed actions. This is what we would expect if LOTC represents the actual observed actions, because the failed actions were incomplete. This fits well with the proposal that LOTC contains an “action recognition” pathway that integrates perceptual information of observed actions into conceptual action representation (Wurm & Caramazza, 2022).

As already noted, the term “goal” can be used to refer to ever finer-grained levels of an action. For example, when observing or executing a grasping action, we could describe the goal as moving the fingers by some parameters so as to enclose them around the object. However, in practice, we would not think of the articulation of the action’s motor components as the goal. By the term “goal”, we are referring to a level of description that generalizes over particular motor specifications and which we can think of as a cognitive level description of an agent’s desired result. In the hierarchical organization of action goals, the desire of the agent can be understood at many levels (Hamilton & Grafton, 2006; Keele et al., 1990). When someone extends their arm toward a bottle of water, the goal can be interpreted as “to grab the bottle”, “to drink”, or “to quench thirst”. An action with a specific immediate goal (“to grab the bottle”) may be followed by upcoming, yet-to-be-observed steps with different immediate goals (e.g., “to drink”). And these immediate goals may be encompassed by a common task goal (“to quench thirst”). Here, our task demand and stimuli design probed only the inference of the immediate goal that describes the physical consequence of the observed action (e.g., “to grab the bottle”), and there is little demand for inferring the mental state of the agent. Further studies are required to uncover where higher-level goals that have stronger demands for mentalizing are represented in the brain independently of outcome.

## Methods

### Participants

Thirty-eight participants participated in the study for cash compensation or course credits. All participants are right-handed and had normal or corrected to normal vision. Six participants were excluded from analysis due to falling asleep in more than two runs or exiting the experiment early. 32 participants (20 female, age from 19 to 36, M = 24.69, SD = 4.78) were included in the final analysis.

### Stimuli

The stimuli set consists of 6-second long videos. Each video depicts one of four possible action goals, balanced across two action types and two object types. They are: “to open a bottle”, “to close a bottle”, “to open a zipper bag” and “to close a zipper bag”. In bottle actions, the agent attempted to pull out or put in the cork. In zipper bag actions, the agent attempted to open or close the bag by pulling the zipper. Each unique goal can end in two different outcomes (success/failure), resulting in 8 unique action events of interests.

To increase the perceptual variance in the stimuli, we used two different exemplars for both object types, filmed the videos from two different viewpoints (from the front or the side of the performer), filmed grasping the zipper or the cork both with right hand and with left hand, and finally flipped every video horizontally. In total, there are 128 videos, balanced across 8 experimental conditions (2 action goals x 2 outcomes x 2 objects). Each condition consists of 16 different perceptual variants.

We also filmed 128 videos where the actions are neither opening or closing involving the same objects as in our main stimuli. For example, the action could be “moving the bottle to one side”, “lifting the bag by its zipper”, “inspecting the contents of the bag”, etc. These videos were presented in catch trials.

### Experiment Design

We used Matlab and Psychtoolbox 3 (Kleiner et al., 2007) for controlling stimuli presentation and recording responses. The fMRI scanning session consists of one T1-weighted structural scan and eight functional task runs.

In the functional runs, the stimuli were presented in an event-related design. Each run started with a 10s fixation phase and ended with 16s of an empty grey background. Each run consisted of 3 blocks, with 12s of fixation between blocks. Each block included 16 trials of interest and 2 catch trials. Each trial consisted of a 6s video and a 2s inter-stimulus interval. The videos were presented in 600 x 450 pixels at the center of the screen. There were 8 functional runs in total, each lasting 472s. In total, there were 384 trials of interest. Across the entire session, each main condition was repeated 48 times, and each unique video was repeated three times.

### Task

To keep participants attentive when watching the videos, we instructed them to press a button using a response box on certain trials. Before entering the scanner, participants were given instructions about the task and went through a practice run on a laptop. They were first shown a subset of 16 opening or closing actions, selected from the 128 videos of interest and balanced across outcomes, objects, and object exemplars. They were asked to carefully watch these videos. Then, they practiced the task they were going to do in the scanner. They were presented with a run consisting of sixteen trials with opening and closing actions and three catch trials that depict neither opening nor closing actions. Participants were instructed to press a button when they recognize an action that was very different from the initial set of videos they watched before. While they were not explicitly told to press a button when they saw an action that is neither opening nor closing, they were able to comprehend the task without issue.

### Data Acquisition

Structural and functional images were acquired using a 3T Siemens MAGNETOM Prisma scanner with a 32-channel head coil. Functional scans were collected using a multi-band echo-planar imaging sequence (EPI) sequence, multi-band factor = 3, repetition time (TR) = 1500ms, echo time (TE) = 28ms, flip angle (FA) = 70°, 45 interleaved slices in a direction from posterior to anterior, field of view (FOV) = 200 mm, slice thickness = 3 mm, voxel size = 3.0 mm x 3.0 mm x 3.0 mm. Each of the eight functional runs comprising the main experiment lasts 7min 51s, consisting of 314 volumes. A high-resolution T1-weighted anatomical scan was collected with an MPRAGE sequence for each participant, consisting of 176 sagittal slices (TR = 2530ms, FA = 7°, slice thickness = 1 mm, voxel size = 1.0 mm x 1.0 mm x 1.0 mm).

### Preprocessing

Functional and structural data were preprocessed in Matlab using Statistical Parametric Mapping Toolbox (SPM12, https://www.fil.ion.ucl.ac.uk/spm/). For each participant, functional images underwent slice-timing correction and then were realigned to the mean functional image. The T1 scan was then coregistered to the realigned mean functional image and then segmented. Functional and structural images were then normalized to the standardized Montreal Neurological Institute (MNI) space. For univariate analysis, functional images were spatially smoothed with an 8mm full width at half maximum (FWHM) Gaussian kernel. For MVPA, the functional scans were smoothed with a 3mm FWHM kernel.

### MVPA Searchlight Analysis

In the first-level analysis, a general linear model was estimated for each participant. The design matrix contained eight predictors of interest (one for each of the eight action conditions), one regressor for catch trials, and six nuisance regressors for translational and rotational head motion parameters. The regressors except for those of head motion parameters were convolved with a canonical hemodynamic response function. A high-pass filter of 128s and an autocorrection model of AR(1) were applied. One beta map was computed per condition per run, resulting in 8 beta maps for each of the eight action conditions (2 goals x 2 objects x 2 outcomes). MVPA classifications were performed on the beta maps.

We performed a searchlight-based classification for each participant in their individual volume space using 12mm searchlight spheres. The classification was implemented with a leave-one-run-out procedure and a linear support vector machine (SVM) classifier, as implemented in CoSMoMVPA Toolbox (https://www.cosmomvpa.org) in Matlab.

First, to identify regions that encode information about actions, we performed a within-object classification of the successful actions. We trained the classifier to discriminate “opening a bottle successfully” vs “closing a bottle successfully” using the betas from all runs except one and tested its accuracy at discriminating these two actions using the betas from the left-out run. This was performed in 8 iterations, using all possible combinations of training runs and testing run. The same was done for actions with zipper bags. The mean accuracy across iterations was computed for each searchlight and was assigned to the voxel at the center of the sphere. To identify voxels that had above-chance accuracies, we entered the individual accuracy maps into a one-tail one-sample t-test. The statistical map was then entered into Monte Carlo Cluster correction to correct for multiple comparisons. The resulting corrected map was used as a mask to constrain the search space of the following searchlight analyses.

To look for regions that encode information related to action goals when the desired outcomes were not achieved, we performed a within-object classification of the failed actions. Using a similar procedure as detailed above, we trained the classifier to discriminate between “failed to open a bottle” vs “failed to close a bottle” and tested on “failed to open a bottle” vs “failed to close a bottle”.

To identify regions that encode information about action goals independently outcome, we performed cross-outcome classifications. We trained the classifier to discriminate between “opening a bottle successfully” vs “closing a bottle successfully” and tested on “failed to open a bottle” vs “failed to close a bottle”. The procedure was repeated for failed bottle actions → successful bottle actions, for all possible iterations and for zipper bag. The mean accuracy of all these combinations were computed.

Finally, to explore whether somewhere in the brain action goals are represented independently of both outcome and of object type, we performed cross-outcome cross-object classifications. We trained the classifier to discriminate between “opening a bottle successfully” vs “closing a bottle successfully” and tested on “failed to open a bag” vs “failed to close a bag”. The procedure was repeated for all possible iterations and combinations and then the mean accuracy were computed.

For each decoding test, the individual accuracies maps from all participants were entered into a one-tailed one-sample t-test against the decoding accuracy expected by chance (50%). To correct for multiple comparisons, the statistical maps were entered into Monte Carlo Cluster correction (10000 iterations) using an initial threshold of p<0.001.

### ROI Definition & Analysis

To examine the action observation network more closely, we selected three bilateral regions of interests based a meta-analysis on action observation (Caspers et al., 2010). We focus on three core regions of the network: ventral premotor cortex (PMv), inferior parietal lobe (IPL), and lateral occipitotemporal cortex (LOTC). We defined the ROIs in each hemisphere by drawing a 12mm sphere centered at the coordinates from the meta analysis (MNI coordinates: left PMv [-50 9 30], right PMv [52 12 26], left IPL [-60 -24 36], right IPL [44 -34 44], left LOTC [-46 -72 2], right LOTC [52 -64 0]).

For each ROI, each participant, and each decoding scheme, we extracted the accuracies for the voxels within the ROI from the searchlight maps and computed the mean accuracy. Then for each ROI and decoding scheme, the mean accuracies from all participants were entered into a one-tailed one-sample t-test against chance (50%). Finally, the p-values were corrected for multiple-comparisons using the FDR method.

## Notes

### Competing Interest Statement

The authors have declared no competing interest.

